# Quantifying Intratumor Heterogeneity by Key Genes Selected using Concrete Autoencoder

**DOI:** 10.1101/2021.09.06.459161

**Authors:** Raihanul Bari Tanvir, Abdullah Al Mamun, Masrur Sobhan, Ananda Mohan Mondal

## Abstract

The tumor cell population in cancer tissue has distinct molecular characteristics and exhibits different phenotypes, thus, resulting in different subpopulations. This phenomenon is known as Intratumor Heterogeneity (ITH), a major contributor to drug resistance, poor prognosis, etc. Therefore, quantifying the levels of ITH in cancer patients is essential, and many algorithms do so in different ways, using different types of omics data. DEPTH (Deviating gene Expression Profiling Tumor Heterogeneity) is the latest algorithm that uses transcriptomic data to evaluate the ITH score. It shows promising performance, has strong similarity with six other algorithms and has an advantage over two algorithms that uses the same type of data (tITH, sITH). However, it has a major drawback since it uses expression values of all the genes (∼20K genes) in quantifying ITH levels. We hypothesize that a subset of key genes is sufficient to quantify the ITH level. To prove our hypothesis, we developed a deep learning-based computational framework using unsupervised Concrete Autoencoder (CAE) to select a set of cancer-specific key genes that can be used to evaluate the ITH score. For the experiment, we used gene expression profile data of tumor cohorts of breast, kidney, and lung cancer from the TCGA repository. Using multi-run CAE, we selected three sets of key genes, each set related to breast, kidney, and lung tumor cohorts. For the three cancers stated and three molecular subtypes of lung cancer, we calculated the ITH level using all genes and key genes selected by CAE and performed a side-by-side comparison. We could reach similar conclusions for survival and prognostic outcomes based on ITH scores derived from all genes and the sets of key genes. Additionally, for subtypes of lung cancer, the comparative distribution of ITH scores derived from all and key genes remains similar. Based on these observations, it can be stated that a subset of key genes, instead of all genes, is sufficient for ITH quantification. Our results also showed that many key genes are prognostically significant, which can be used as possible therapeutic targets.

## 1 Introduction

Intratumor Heterogeneity (ITH) refers to different types of tumor cell subpopulations within a tumor [1]. Even though these cell subpopulations have the same origin (tumor tissue, patient), they exhibit different phenotypes and possess different molecular characteristics. ITH is one of the main challenges for targeted cancer therapy, as the difference in tumor cells and their microenvironments makes it harder for targeted cancer therapy to eradicate cancer cells [2][3]. Therefore, an accurate assessment of ITH is essential to understand the tumor dynamics and development of effective and durable therapeutic strategies. The causes of ITH can be many depending on different levels, such as genome, epigenome, transcriptome, etc. [4]. For example, reduced DNA damage mechanisms, microenvironmental factors (hypoxia, acidosis, etc.) [5], subclonal evolution [2], etc. are contributors of ITH in genomic level. Methylation of tumor suppressor genes is an example of ITH at epigenomic level [6]. Different patterns of gene expression contribute to ITH at transcriptome level, which is observed to mirror ITH in genomic or epigenomic level or both [5][7]. This makes transcriptomic data suitable for quantifying ITH.

There are different algorithms for quantifying ITH, such as ABSOLUTE [8], MATH [9], EXPANDS [10], PhyloWGS [11]. These algorithms use genomic data, such as – copy number alterations (CNA), somatic mutation profiles, etc. There are also algorithms that take advantage of transcriptome profile that mirrors ITH in genomic and/or epigenomic level, such as-tITH [12], sITH [13], and DEPTH [14]. Among the algorithms that quantify ITH, DEPTH showed strong similarity in calculating a degree of ITH with other genome-based and transcriptome-based methods. DEPTH has certain advantages over its two other transcriptome-based counterparts, tITH and sITH. First, unlike the other two, DEPTH can calculate ITH score without normal samples. Second, it calculates ITH score using only gene expression data. tITH requires protein-protein interaction (PPI) network along with gene expression data.

DEPTH algorithm also has certain drawbacks. While it uses only gene expression data, it uses expression values of all genes (∼20K genes) in calculating ITH score. While calculating ITH, not all the genes may contribute to ITH. Additionally, not all genes may have alterations in gene expression patterns due to the effect of ITH in genomic and/or epigenomic level. There may be a subset of genes which is sufficient in calculating ITH score at transcriptome level. In this study, we propose a deep learning-based computational framework based on unsupervised concrete autoencoder (CAE) [15] to identify a set of key genes that can be used to quantify ITH. CAE can select the most informative subset of features from a large set of features which also have a minimum reconstruction error. First, we select a set of key genes from breast (BRCA), kidney (KIRC) and lung cancer (LUAD) using the expression profile data from TCGA repository. Then we calculate the ITH score for the tumor patients in three cancer types using all genes and the key genes employing DEPTH algorithm. We also calculate DEPTH score using all genes and key genes for different molecular subtype of LUAD. In both cases, we show that a set of key genes (k=100) can achieve similar results as using all genes (k=20,000+). Our contribution is the development of a computational framework to identify key genes that can be used in evaluating the ITH score. In other words, we do not need all the genes used in the previous work in evaluating the ITH score.

## 2 Materials and Methods

### 2.1 Dataset Collection

We collected gene expression dataset of TCGA Breast Carcinoma (BRCA), Lung Adenocarcinoma (LUAD) and Kidney Renal Carcinoma (KIRC) from USCS Xena Browser database [16]. Each dataset contains expression profiles of 20,530 mRNAs. The distribution of tumor and normal samples for three cancers is given in Table 1.

**Table 1:** Summary of gene expression data of BRCA, LUAD and KIRC cohorts from TCGA.

| Cancer Type | # Tumor | # Normal | # Genes |
| --- | --- | --- | --- |
| BRCA | 1097 | 114 | 20530 |
| KIRC | 533 | 72 | 20530 |
| LUAD | 515 | 59 | 20530 |

### 2.2 Concrete Autoencoder to Select Cancer-specific Key Genes

Concrete Autoencoder (CAE) [15], an unsupervised deep learning approach, is used to identify cancer-specific key genes. CAE identifies features that are most informative for a given dataset [15], [17].

#### CAE Architecture and Working Principle

Figure 1 shows the general architecture of CAE. CAE differs from the standard Autoencoder in the encoder part, where CAE employs a concrete selector layer. This selector layer is based on Concrete distribution [18] or Gumbel Softmax distribution [19], which is a relaxed variant of discrete distribution. The selector layer is used to incorporate discrete distribution into deep learning algorithms, and as an example, CAE uses it to learn a subset of features that are most informative and produces minimum reconstruction error. In the learning phase, the selector layer learns a subset of features, which depends on a hyperparameter called Temperature (T), which is gradually lowered during training phase to a low value using a simple annealing scheduling. This gradual decrease in temperature helps the concrete distribution to learn and select a definite subset of features [15]. Unlike the encoder part of the CAE, the decoder part resembles closely with the standard Autoencoder.

**Figure 1:**
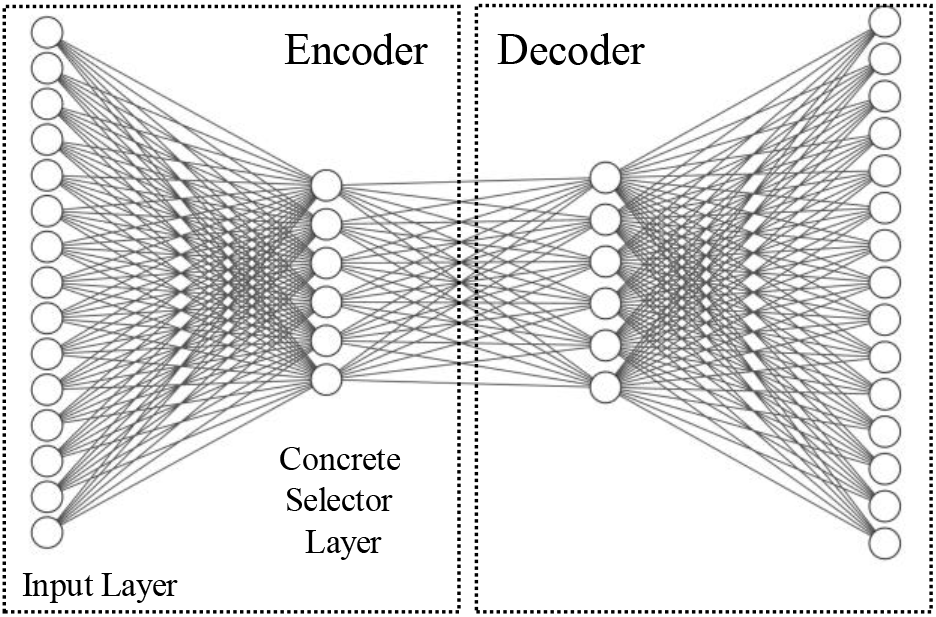
The general architecture of CAE. The encoder part contains the input layer and the concrete selector layer. The decoder in this figure is a 2-layer neural network, with the final layer having the same number of nodes as the input layer.

In the selector layer, each unit selects a unique feature with the highest probability from the original feature space. Thus, CAE selects the most informative subset of features and the reconstruction of original feature space using the selected subset of features produces minimum reconstruction error. In the original Autoencoder, the features learned at the encoder part are latent features, whereas the features learned at CAE are actual features.

CAE was trained on the each of the gene expression data of BRCA, KIRC and LUAD and 100 features were selected in each run. While training CAE, the dataset was divided randomly into 80/20 split for training and testing.

### 2.3 Selection of Hyperparameter and Tuning

We kept three of the parameters the same as used in the original CAE developed by Abid et al. [15]. These parameters are (i) leaky ReLU as activation function with a threshold value of 0.1, (ii) 10% dropout rate, and (iii) temperature T reduced using a simple annealing schedule from 10 to 0.1, For decoder part of CAE, we used a neural network with two hidden layers, each layer containing 300 units as used in our previous work with expression profile data [17].

Number of epochs and learning rate was tuned using random search [20]. For *<u>tuning learning rate</u>*, we varied the number of epochs from 2000 to 10000. The number of epochs were set to high since the number of samples are low (ranges between 515 for KIRC to 1097 for BRCA) compared to the feature space of ∼20K genes. Another reason for using large number of epochs is to reach the high mean-max probability. Mean-max probability is the average of maximum probability value for each feature selected by a node in the concrete selector layer. Each node in concrete selector layer selects the feature with the highest probability. Therefore, one of the main goals in training of CAE is to make sure that the mean-max probability is high (≥ 0.98). The learning rate was varied from 0.001 to 0.01 with a step size of 0.001 and it was found that 0.002 was optimum in reaching the mean-max probability ≥ 0.98. The number of epochs set for BRCA, KIRC and LUAD were 3500, 9000, and 8000, respectively. The average number of epochs to reach the threshold of mean-max probability ≥ 0.98 were 3066, 8191 and 7250 for BRCA, KIRC, and LUAD datasets, respectively.

### 2.4 Training CAE

Figure 2 shows the characteristics curve for CAE or an instance of the training behaviors of CAE for LUAD dataset. The hyperparameter, Temperature (T) was reduced using a simple annealing schedule from 10 to 0.1 from the start epoch to the last.

**Figure 2:**
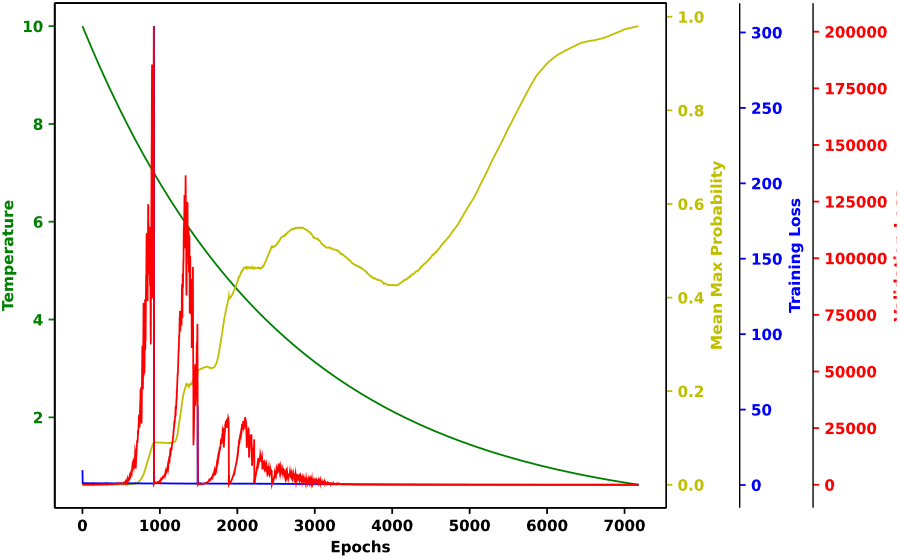
The characteristics curve of CAE. Reconstruction errors with training data (blue) and validation data (red) were plotted. The green and yellow curve represent the values of Temperature (T) and mean-max probability, respectively.

The reconstruction errors (loss) for training set and validation set are plotted using blue and red curve respectively. It shows that both errors were relatively high during the early phase of training as expected and both reach to a minimum plateau at the end. Also, the mean-max probability finally approaches 1.0 (yellow curve).

#### Implementation

The CAE was implemented using Keras (https://keras.io/). Experiments were conducted in parallel manner on high performance cluster with NVIDIA Quatro K620 GPU with 384 cores and 2GB memory devices.

### 2.5 ITH scoring method

To calculate the score of Intratumor Heterogeneity (ITH), we used a scoring method named-Deviating Gene Expression Profiling Tumor Heterogeneity, or DEPTH in short [14]. This method is mRNA-based, and it only uses the gene expression values of tumor samples and normal samples. The DEPTH score of a tumor sample TS is defined as –

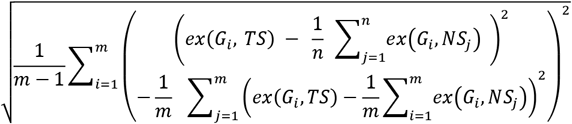

Where, TS is the tumor sample for which the score is being calculated. *G*_*i*_ is the i-th Gene and *m* is the number of genes. *n* is the number of normal patients. *ex*(*G*_*i*_, *S*) expression of gene *G*_*i*_ in sample *S*. It assigns a score to each patient. It is based on standard deviations of the gene expression values variations in tumors from mean value of normal samples. We calculated ITH score for each of the cancer patients of BRCA, KIRC and LUAD employing DEPTH using two sets of genes. One score using all the genes as used in [14] and the other using only the key genes selected by multi-run CAE.

### 2.6 Survival Analysis

Survival Analysis was performed to check whether two groups of patients based on high and low ITH scores are significantly distinguishable in prognosis. In our analysis, the event of interest is the death of cancer patients.

#### Survival Analysis Based on ITH Scores

Samples were sorted in descending order of ITH score, and then the top one-third and the bottom one-third of the total samples were taken as two groups. This analysis was done to compare the prognostic importance of ITH scores derived using all genes [14] and key genes (our study) (see Section III-C, III-D).

#### Survival Analysis Based on Each of Key Genes

The cohort was divided into two groups based on median of the expression values of the gene in question. This survival analysis was done to check how many of key genes are prognostically important.

In both cases, after forming two groups, Kaplan-Meier curve was plotted, and Log-rank test was performed to check the statistical significance of difference in survival function.

## 3 Results and Discussion

### 3.1 Selecting Key Genes – Multi-Run CAE

Due to the stochastic nature of CAE, the model was trained 10 times, and in each run, 100 features were selected for each cancer cohort - BRCA, KIRC and LUAD. Figure 3(a) shows the stochastic nature of CAE since only 16 features are common between three single-run CAE. In case of 10-run CAE, top 100 features were selected from the combined list sorted in descending order based on the frequency of appearance of a feature in 10 runs. It is clear from Figure 3(b) that there are 53 genes common between three batches of 10-runs. Thus, multi-run approach was adopted to select the robust set of features. To select the key features, the top 100 frequent features were chosen based on the assumption that the more frequent a feature in different runs, the more informative the feature is. The summary of 10-run CAE for the three cancer cohorts are shown in Table 2. The combined lists of features derived from 10-run CAE consist of 469, 527, and 435 genes for BRCA, KIRC, and LUAD, respectively. The range of frequency of top 100 features is 3 to 10 for each cancer cohort, which means that the most frequent features appeared in all 10 runs and the least frequent one appeared in 3 runs.

**Figure 3:**
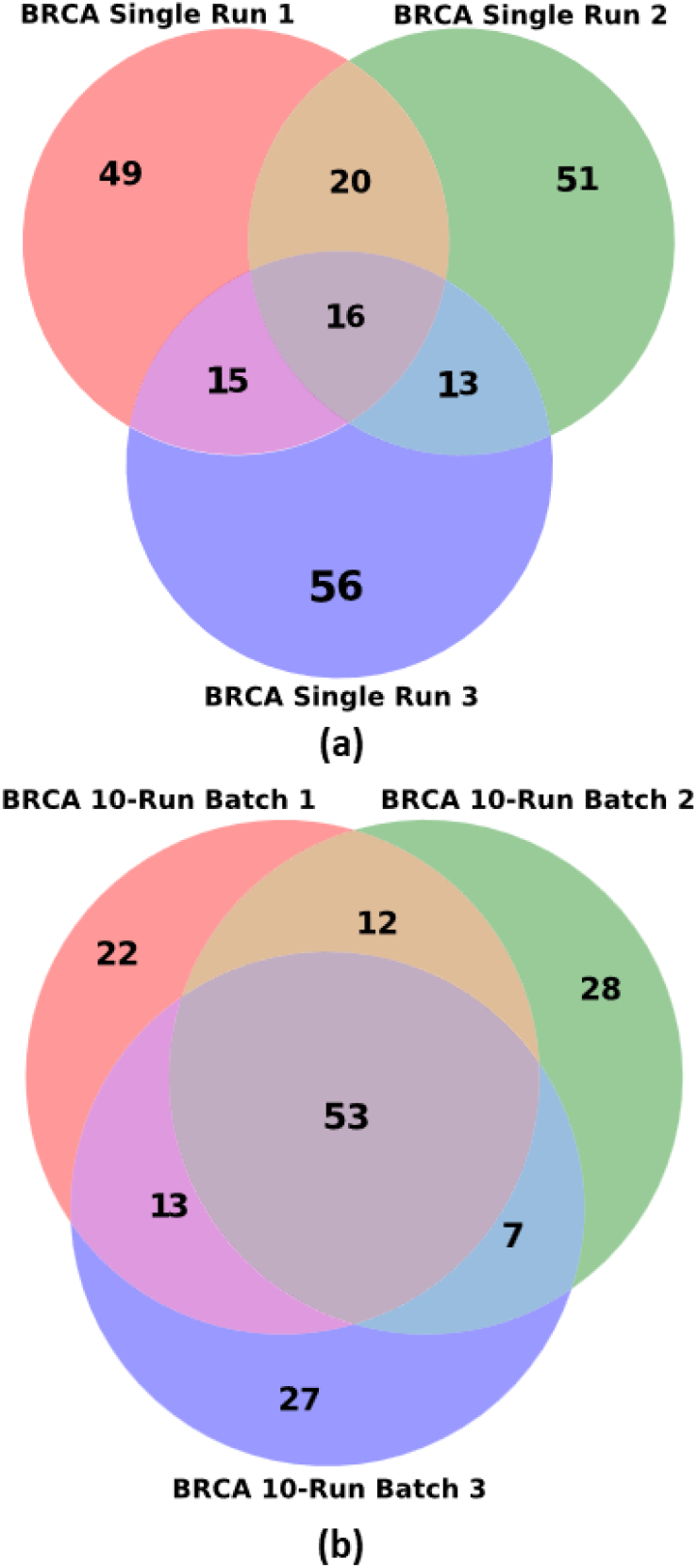
Selecting robust set of features. (a) Venn diagram of three sets of 100 genes from three single-run CAE; CAE produces only 16 features common between three single runs. (b) three sets of top 100 features from 10-run CAE. 10-run CAE produces more features (53 genes) common between three batches of runs. Thus, multi-run CAE produces robust set of features.

**Table 2:** Summary of Multi-run CAE runs for BRCA, KIRC, LUAD cohort.

| Cancer Type | # Genes in Multi-run | Frequency Range in Top 100 |
| --- | --- | --- |
| BRCA | 469 | 3-10 |
| KIRC | 527 |  |
| LUAD | 435 |  |

### 3.2 Commonality Check

We investigated the commonality of the key gene sets of three cancers, which is shown by the Venn diagram in Figure 4. It shows that there is no common gene between three gene sets. But there are a few genes common between each pair of gene sets: 5 between BRCA and LUAD, 3 between KIRC and BRCA, and 3 between KIRC and LUAD. Since the size of each set is 100 and there are only few genes common between two sets and none between the three sets, thus, the key gene sets are cancer specific.

**Figure 4:**
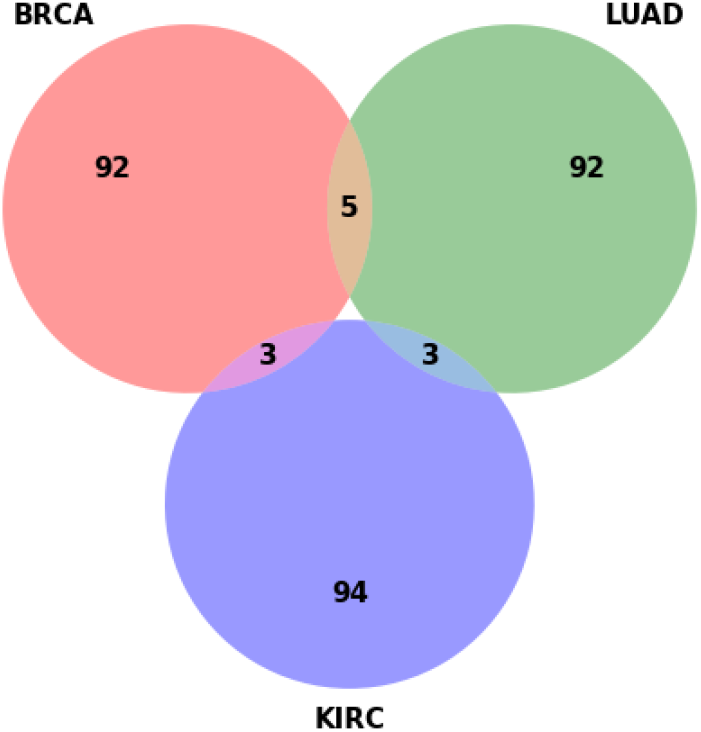
Venn diagram of three sets of Key genes for BRCA, KIRC and LUAD cohort.

### 3.3 All Genes vs. Key Genes in ITH scoring: Case - Whole Cancer Cohort

In this section, we compare the ITH scores calculated for BRCA, KIRC and LUAD cohorts using two different sets of genes: (i) DEPTH score calculated using all genes [14] and (ii) DEPTH score calculated using only the key genes selected by the multi-run CAE system (our work). Survival analysis is used to compare the two TH scores. Figure 5 presents the results of survival analyses, Kaplan Meier plots, for three cancer cohorts - BRCA (a-b), LUAD (c-d), and KIRC (e-f) - based on ITH scores derived from all and key genes. It is evident from Kaplan Meier plots that high DEPTH scores are related to poor prognosis, and low DEPTH scores have higher chance of survival. By comparing plots (a) with (b), (c) with (d), and (e) with (f), it is clear that the similar results can be achieved for each cancer cohort by using only the 100 key genes. Thus, we do not need all 20,000 genes to evaluate the ITH scores as done in [14].

**Figure 5:**
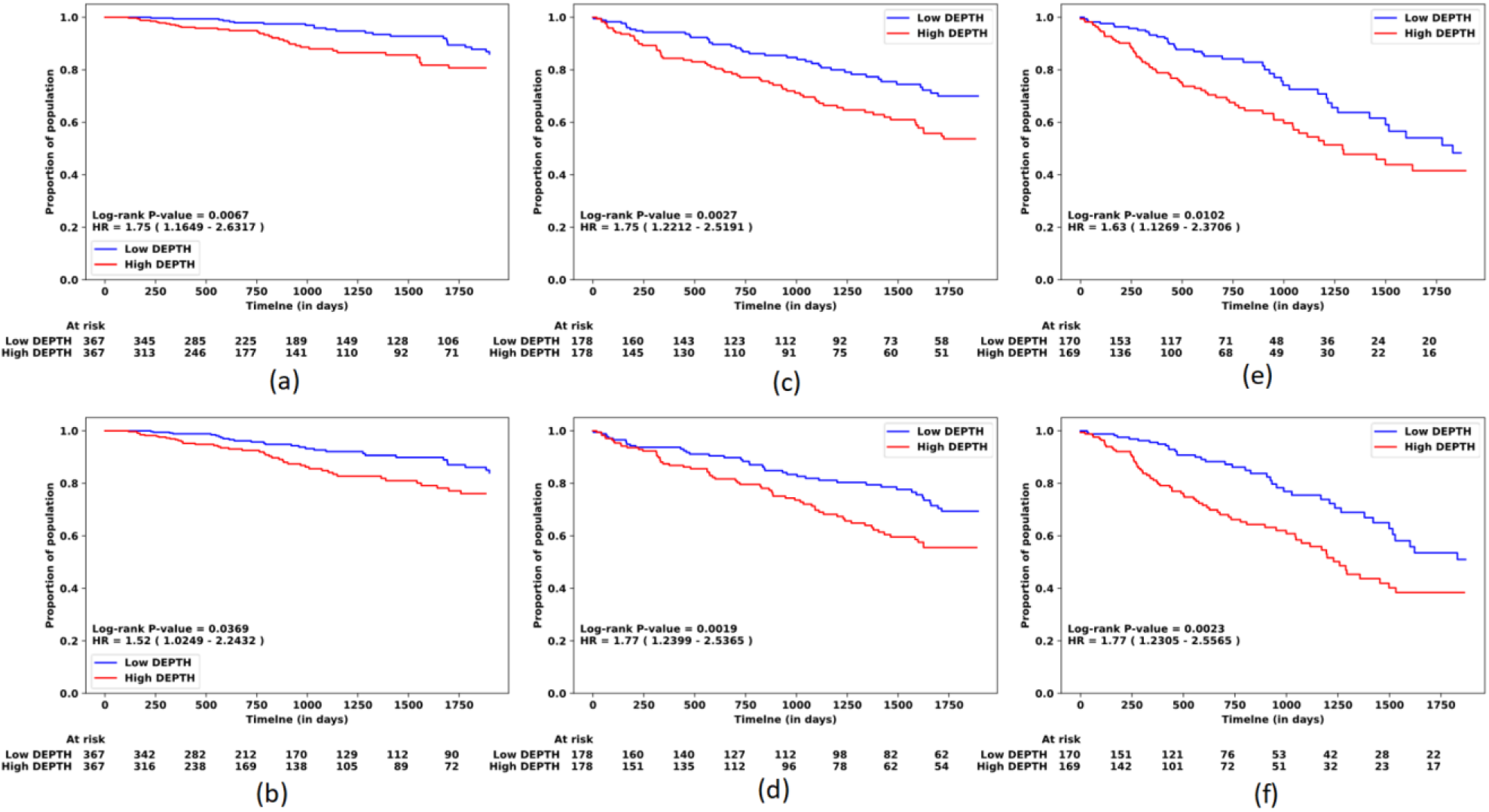
Prognostic analysis using DEPTH scores. Kaplan Meier plots for BRCA(a)(b), KIRC(c)(d), and LUAD(e)(f). Top-row KM plots ((a),(c),(e)).: DEPTH scores calculated using all genes. Bottom-row KM plots ((b),(d),(f)): DEPTH scores calculated using key genes.

The hazard ratios and the log-rank P-values are comparable though there are slight differences between two corresponding plots. For example, for BRCA, P-value of 0.0067 and Hazard Ratio of 1.75 were found in survival analysis using all genes, whereas key genes produced a P-value of 0.0369 and Hazard Ratio of 1.52. But unlike BRCA, KIRC and LUAD show better outcome for key genes than all genes by producing reduced P-values and increased hazard ratios. It is evident that, only the key genes can be used to quantify ITH of each patient in the whole cohort of particular cancer type.

Then, we compared the low DEPTH groups of Figure 5(a) (blue curve) and 5(b) (blue curve) from BRCA. Comparing the two sample sets from two low-DEPTH groups, each containing 367 patients, we found 283 (77%) are common. When we did survival analysis again using these two groups we found the log-rank test p-value was 0.263 and the hazard ratio was 0.78, as shown in Figure 6(a). This observation suggests that the low-DEPTH groups from figures 5(a) and 5(b) are not significantly distinguishable. Similar results were also found for high DEPTH groups from Figures 5(a) and 5(b) (red curves) with log-rank test p-value 0.7397 and hazard ratio 0.94, which means the two groups of patients are statistically indistinguishable (Figure 6(b)). It is evident from these analyses that the blue groups from Figures 5(a) and 5(b) and red groups from Figures 5(a) and 5(b) are not statistically different, thereby somewhat similar. Similar trends were also observed for LUAD and KIRC.

**Figure 6:**
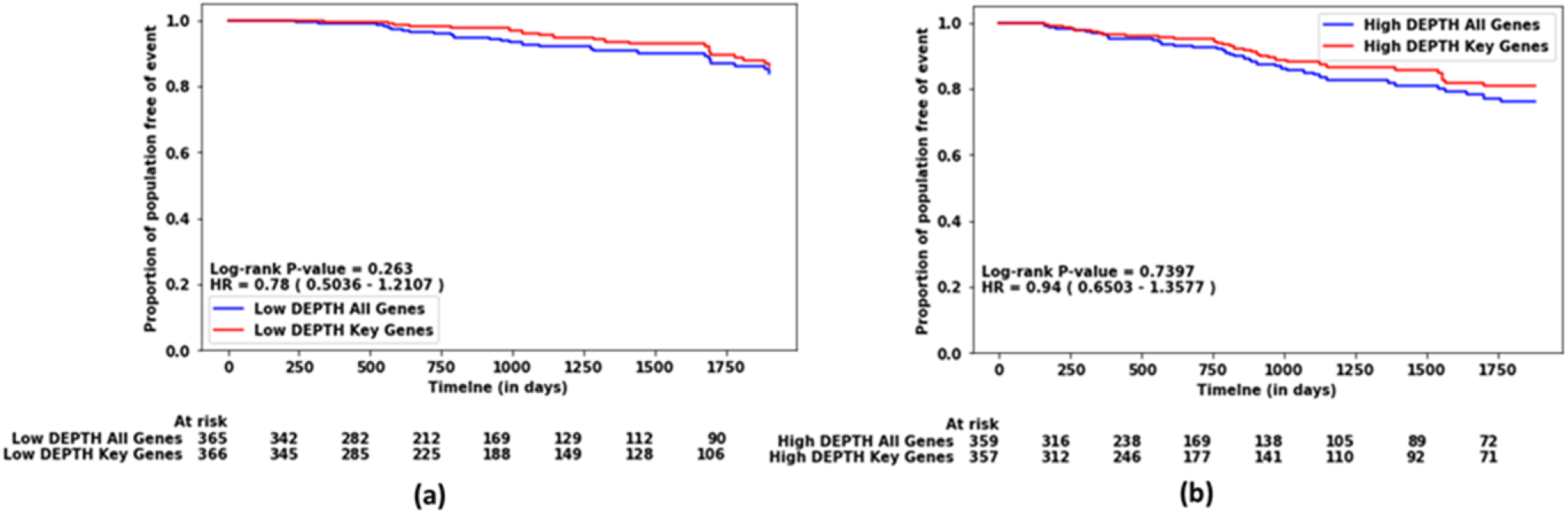
Comparison of two groups of patients with two low-DEPTH scores and two high-DEPTH scores for BRCA cohort. (a) Survival curve of two groups with low-DEPTH scores using all genes and key genes, (b) Survival curve of two groups with high-DEPTH scores using all genes and key genes. score are selected by selecting the patients with bottom one-third of the DEPTH scores. Similarly, patients with high DEPTH score are selected by selecting the patients with top one-third of the DEPTH scores.

### 3.4 All Genes vs. Key Genes in ITH scoring: Case - LUAD subtypes

In this section, we show the comparison of ITH scores (DEPTH) for LUAD subtypes calculated using all genes versus key genes. Of 435 LUAD patients, 55, 34 and 54 are labeled as Terminal Respiratory Unit (TRU), Proximal Proliferation (PP) and Proximal Inflammation (PI), respectively. This molecular subtyping was done in [21]. We compare ITH score for two cases: all genes vs. key genes. Figure 7(a) shows the prognostic difference between TRU and PI and PP combined, from where it is evident that, TRU subtype is prognostically favorable, and it has higher chance of survival than other two subtypes combined. Figures 7(b) shows the ITH score distribution for three subtypes, using all genes and key genes. From Figure 7(b), it is seen that the subtype TRU has comparatively lower values in ITH score than other subtypes, which results in higher chance of survival for TRU subtype. The ITH scores for a patient derived using all and key genes were different. So, min-max normalization on whole LUAD cohort was performed to bring the DEPTH score distribution in the same scale. The overall comparative distribution remained the same (Fig. 7(b)).

**Figure 7:**
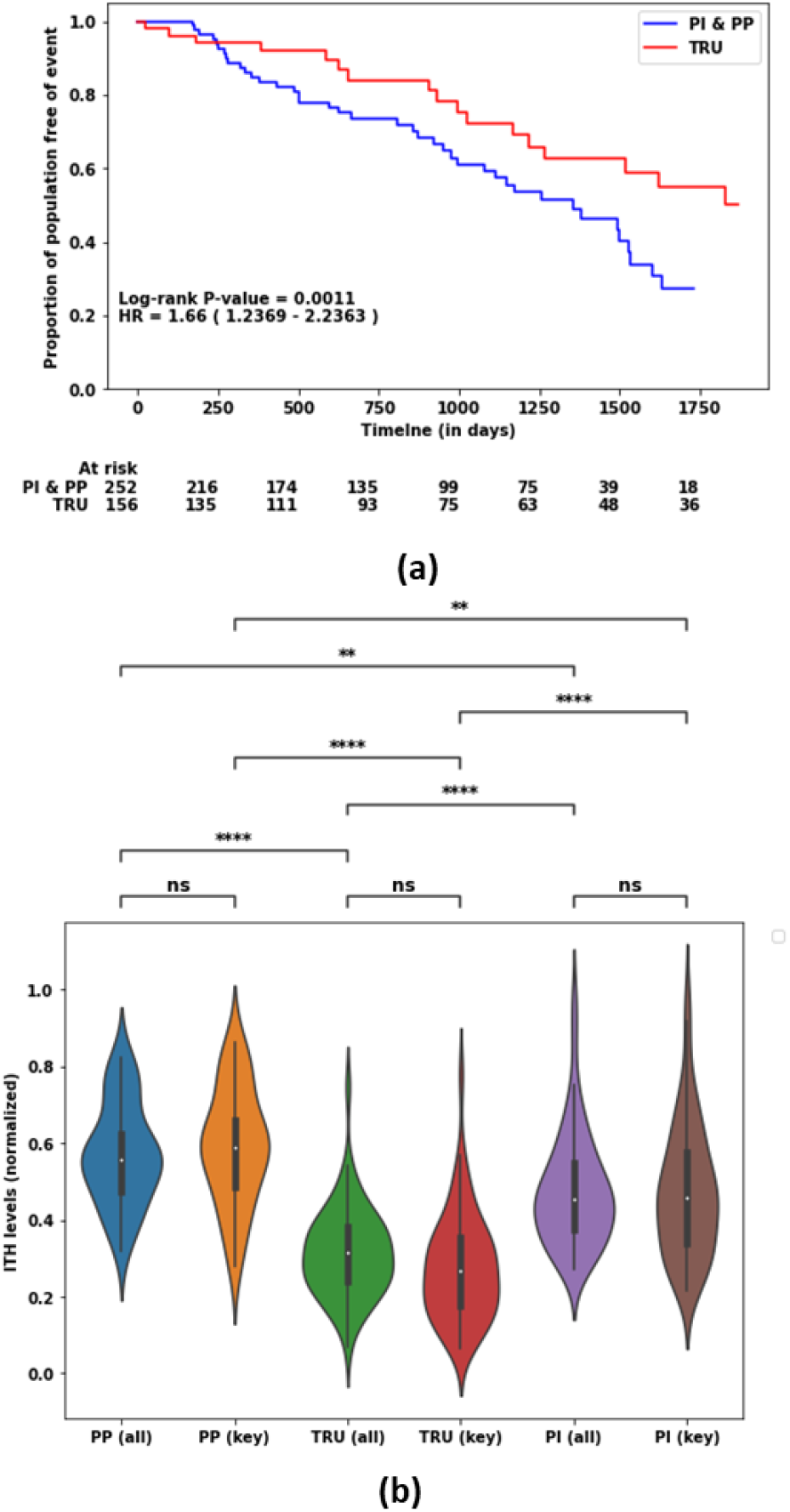
Comparison of ITH scores using all genes (labeled as ‘all’) and key genes (labeled as ‘key’) in molecular Subtypes of LUAD. (a) Kaplan-Meier Curve comparing the survival of TRU subtype vs PI and PP subtypes combined. (b) Distribution of Min-max normalized ITH score (DEPTH) in three molecular subtypes of LUAD in violin plots. The Mann-Whitney-Wilcoxon test between two distributions was performed and the p-value significance is marked by stars. p-value annotation legend: ns (not significant): 0.05 < p ≤ 1, *: 0.01 < p ≤ 0.05, **: 0.001 < p ≤ 0.01, ***: 0.0001 < p ≤ 0.001, ****: p ≤ 0.0001

To compare the distribution of the DEPTH scores using all and key genes, we performed correlation analysis, which is given in Table 3. From Table 3, we see that there is relatively high correlation between DEPTH scores of PP, TRU, and PI subtypes using all and key genes from Figure 7(b) and all the coefficient values were statistically significant. Based on this observation, it can be concluded that to calculate the ITH score, only the key genes will suffice.

**Table 3:** Correlation of DEPTH score distribution calculated using all genes and key genes for three molecular subtypes of LUAD and their p-values. The Pearson and Spearman correlation coefficients and the correaponding p-values are listed.

| Correlation |  | Pearson | Spearman |
| --- | --- | --- | --- |
| PP (all) vs PP (key) | Coefficient | 0.6031 | 0.6091 |
|  | p-value | 0.0001 | 0.0001 |
| TRU (all) vs TRU (key) | Coefficient | 0.7756 | 0.7466 |
| | p-value | $3.57 \times 10^{-12}$ | $5.94 \times 10^{-11}$ |
| PI (all) vs PI (key) | Coefficient | 0.7946 | 0.8078 |
| | p-value | $7.32 \times 10^{-13}$ | $7.32 \times 10^{-13}$ |

### 3.5 Survival Analysis of Key Genes

Survival analysis was performed on each of the key genes from their respective cancer cohort to identify whether they possess prognostic capabilities. Figure 8 shows the forest plot of the prognostically significant genes, along with the summary of survival analyses in terms of Logrank P-value and Hazard Ratio with 95% confidence interval. The thresholds for prognostically significant genes are Logrank P-value **≤** 0.05 and Hazard Ratio, HR ≠ 1. Of 100 key genes for BRCA, 15 were prognostically significant, as shown in the forest plot in Figure 8. Similarly, for KIRC and LUAD, 30 and 61 genes were found to be prognostically significant.

**Figure 8:**
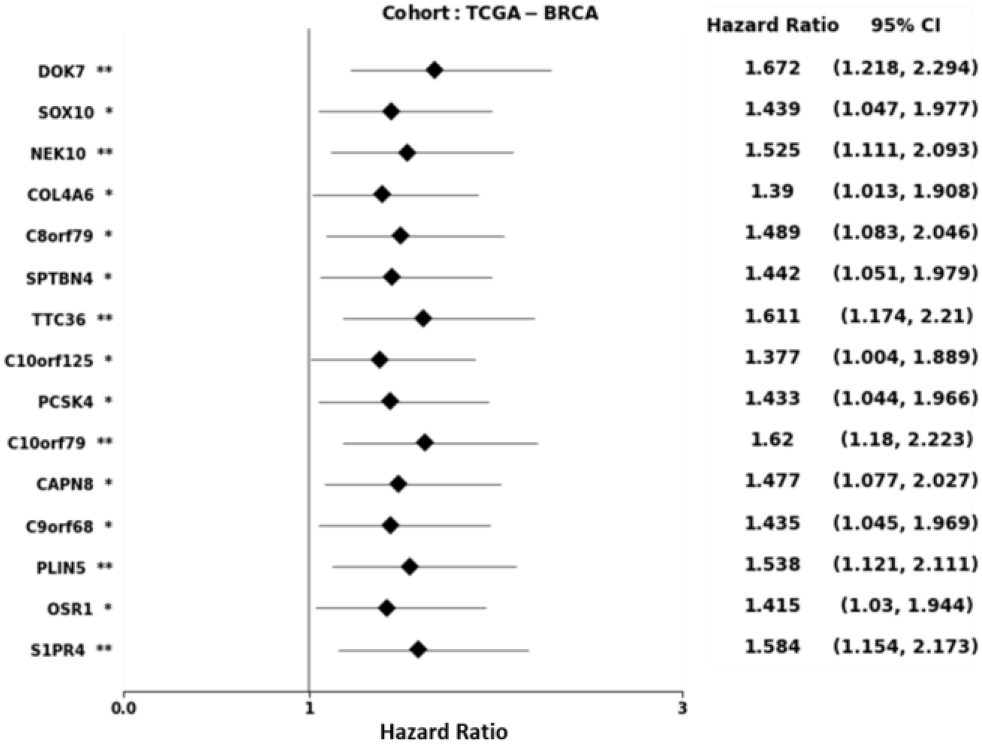
Forest plot of prognostically signifcant key genes for BRCA. It shows the values of Hazard Ratio (diamond shape) with 95% confidence interval (straight line crossing the diamond). The logrank-test p-value is represented by the number of stars. (*: 0.01 < p ≤ 0.05, **: 0.001 < p ≤ 0.01, ***: 0.0001 < p ≤ 0.001).

### 3.6 Conclusion and Future Direction

In this study, we hypothesize that a subset of key genes, instead of all genes (∼20,000 genes), is sufficient to evaluate the ITH scores of individual tumors. To prove our hypothesis, we developed a computational framework based on a multi-run concrete autoencoder that selects a set of key genes for a tumor cohort that can be used to evaluate the ITH score for an individual tumor. It was shown that, instead of all genes used in the existing state-of-the-art method, a subset of key genes is sufficient to evaluate the ITH score using gene expression profile data. We showed that ITH scores derived using the subset of key genes and all genes produced similar survival and prognostic outcomes for three different cancers (BRCA, KIRC, and LUAD) and molecular subtypes of LUAD. This study concludes that a subset of key genes is sufficient to quantify the ITH at the transcriptome level. We also showed that many of these key genes are prognostically significant, which can be investigated further as possible therapeutic targets.

The ITH depends on individual genetic and epigenetic heterogeneity, and transcriptome mirrors both genetic and epigenetic heterogeneity of an individual tumor. Thus, there should exist a patient-specific set of genes whose expression profiles should dictate the ITH of an individual tumor. In the present study, we used the same set of key genes to evaluate the ITH for all patients of a particular tumor type, which is a limitation. In the extended version of this work, we will identify the patient-specific set of key genes to evaluate the ITH.

## ACKNOWLEDGMENTS

This work has been partially supported by the NIH/NIMHD Award #1U54MD012393-03, NSF CAREER Award #1901628, NSF RPAID Award #2037374.

